# Identification of orthotropic material parameters for acute, necrotic, fibrotic and remodelling myocardial infarcts in the rat

**DOI:** 10.1101/754754

**Authors:** Mazin S. Sirry, Laura Dubuis, Neil H. Davies, Jun Liao, Thomas Franz

**Affiliations:** Division of Biomedical Engineering, Department of Human Biology, University of Cape Town, Observatory 7925, South Africa; Stellenbosch Institute for Advanced Study (STIAS), Wallenberg Research Centre at Stellenbosch University, Stellenbosch, South Africa; Department of Biomedical Engineering, University of Medical Sciences and Technology, P.O. Box 12810, Khartoum, Sudan; Cardiovascular Research Unit, Chris Barnard Division of Cardiothoracic Surgery, University of Cape Town, Observatory 7925, South Africa; Department of Bioengineering, University of Texas at Arlington, Arlington, TX 76010, USA; Bioengineering Science Research Group, Engineering Sciences, Faculty of Engineering and the Environment, University of Southampton, Southampton SO171BJ, UK; CSIR Centre for High Performance Computing, Rosebank 7700, South Africa

**Keywords:** Material optimisation, soft tissue, biaxial, inverse methods, anisotropic, genetic algorithms

## Abstract

Finite element (FE) models have been effectively utilized in studying biomechanical aspects of myocardial infarction (MI). Although the rat is a widely used animal model for MI, there is a lack of material parameters based on anisotropic constitutive models for rat myocardial infarcts in literature. This study aimed at employing inverse methods to identify the parameters of an orthotropic constitutive model for myocardial infarcts in the acute, necrotic, fibrotic and remodelling phases utilizing the biaxial mechanical data developed in a previous study. FE model was developed mimicking the setup of the biaxial tensile experiment. The orthotropic case of the generalized Fung constitutive model was utilized to model the material properties of the infarct. The parameters of Fung model were optimized so that the FE solution best fitted the biaxial experimental stress-strain data. A genetic algorithm was used to minimize the objective function. Fung orthotropic material parameters for different infarct stages were identified. The FE model predictions best approximated the experimental data of the 28 days infarct stage with 3.0% mean absolute percentage error. The worst approximation was for the 7 days stage with 3.6% error. This study demonstrated that the experimental biaxial stress-strain data of healing rat infarcts could be successfully approximated using inverse FE methods and genetic algorithms. The material parameters identified in this study will provide an essential platform for FE investigations of biomechanical aspects of MI and the development of therapies.

## 1. Introduction

Computational biomechanics has been effectively utilized for disease investigation and therapy optimisation providing answers to many clinical questions. Among several applications in the cardiovascular area, finite element (FE) models have been employed for studying the mechanical aspects of myocardial infarction (MI) (Wenk et al., 2011) and its emerging treatment through intramyocardial injection of biomaterials (Kortsmit et al., 2013; Miller et al., 2013; Wise et al., 2016).

The passive constitutive behaviour of myocardium has been modelled as transverse-isotropic (Guccione et al., 1991; Humphrey et al., 1990) or orthotropic (Costa et al., 2001; Holzapfel and Ogden, 2009) incompressible finite elastic material using hyperelastic pseudo-strain energy functions. Inverse methods are utilized for estimating the parameters of these constitutive models by fitting FE solution to experimental data (Usyk and McCulloch, 2003). Such data are obtained either from mechanical testing of isolated heart samples (Gupta et al., 1994) or by measuring regional deformation from the whole heart (Fomovsky and Holmes, 2010; Lee et al., 2011; Wenk et al., 2010). Genetic algorithms (GA) are global search methods based on the concept of biological evolution. GA and FE methods have been jointly employed for identification of material parameters of biological tissue (Khalil et al., 2006; Kichula et al., 2014; Wang et al., 2006; Yeoman et al., 2009).

Material parameters for infarcted myocardium are widely available in literature for different animal species. Although the rat is a widely used animal model for MI studies, material parameters for healing rat myocardial infarcts remain underrepresented especially those based on anisotropic constitutive models. This emphasizes the need to identify material parameters of an anisotropic constitutive model for myocardial infarcts in rats.

In our previous work (Sirry et al., 2016a), we experimentally characterised the mechanical properties of rat myocardial infarcted tissue immediately, 7, 14 and 28 days after the infarct induction representing the acute, necrotic, fibrotic and remodelling healing phases of the infarct, respectively. The current study aimed at identifying the parameters of an orthotropic constitutive model for the four healing stages of rat myocardial infarcts utilizing mean biaxial stress-strain data (Sirry et al., 2016b). A biaxial tension inverse FE model was developed mimicking the setup of the experimental biaxial test (Sirry et al., 2016a) and GA were employed to minimize the error between the predicted and experimental data.

## 2. Methods

### 2.1 Biaxial tension finite element model

Abaqus CAE 6.12-2 (Dassault Systèmes, Providence, RI, USA) was employed to build and run a biaxial tension FE model following the experimental setup described in our previous work (Sirry et al., 2016a). A three-dimensional (3D) model of the tissue sample was generated by extruding a square area of length (*l*) by depth (*T*) as illustrated in Figure 1. The endocardial and epicardial surfaces were represented by *x-y* planes at *z*=0 and *z*= *T*, respectively. As a result, the *x* and *y* axes of the model represented the cardiac circumferential and longitudinal axes, respectively. The eight suture needles used in the experiment were modelled using cylindrical transmural partitions, indicated by C1 to C8 in Figure 1 (a), of 0.4 mm diameter (the physical diameter of the needles). The two adjacent cylinders on each side were separated by *l*/3 while all cylinders were positioned 2 mm away from the edge. Four reference points, RP1 to RP4 in Figure 1 (a), were defined at the central area of the epicardial surface of the model representing the four optical markers used in the experiment. A tie constraint was defined to couple the movement of the four reference points to the local movement of the epicardial surface.

**Figure 1:**
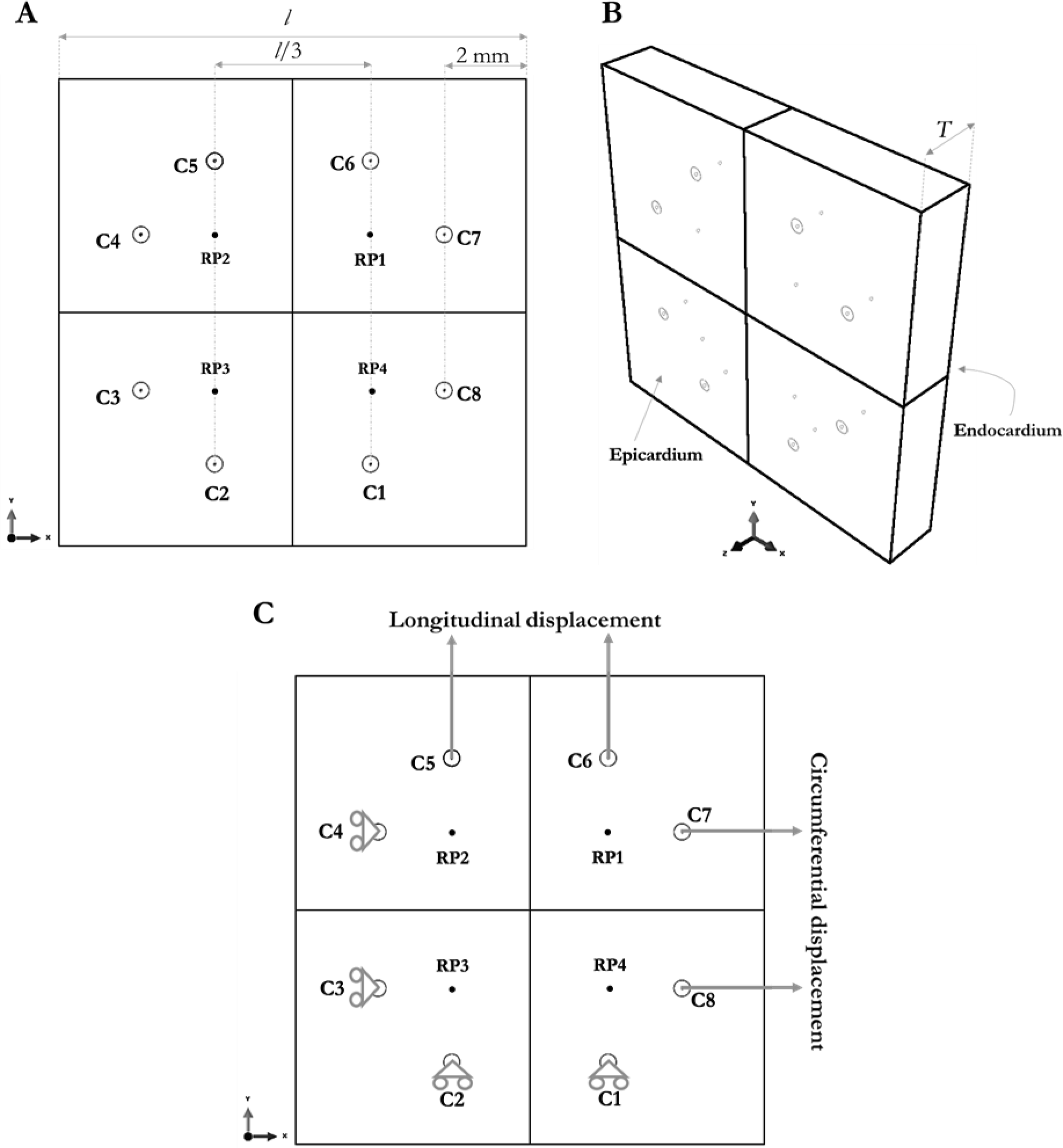
Development of the FE model: (a) Two-dimensional sketch of the FE model showing the positions of suture needles (C1-8) and reference points (RP1-4). (b) Extruded geometry of the specimen showing epicardial and endocardial surfaces. (c) Illustration of the boundary conditions defined in the model.

Four models were developed representing the four infarct groups, i.e. immediate (0 day), 7, 14 and 28 days. The model dimensions were defined based on average physical dimensions measured from samples within each infarct group. The model was discretized using linear hexahedral reduced-integration hybrid elements (C3D8RH). The mesh contained 1896 elements. The orientation of myocardial fibres was incorporated in the model by a user-defined material orientation (ORIENT) subroutine. The fibre angles ranged linearly from −50° at the epicardial layer to 80° at the endocardial layer (Chen et al., 2003).

### 2.2 Boundary conditions

Displacement boundary conditions were applied to the cylinders to restrain the model (C1 to C4) and to apply the tensile loads (C5 to C8) as illustrated in Figure 1 (c). The translational movement of C1 and C2 was constrained in the *y*-axis while the translational movement of C3 and C4 was constrained in the *x*-axis. All cylinders were allowed to rotate freely. Circumferential tensile load was applied by displacing C7 and C8 in the *x*-axis while longitudinal tensile load was applied by displacing C5 and C6 in the *y*-axis.

### 2.3 Constitutive model

From a number of anisotropic hyperelastic constitutive models available in Abaqus, the orthotropic case of the generalized Fung model was utilized to model the mechanical behaviour of the infarct. The generalized Fung strain energy potential is based on the exponential form suggested by Fung et al. (1979) appropriately generalized to arbitrary 3D states following Humphrey (1995). Infarct material was assumed to be fully incompressible. The generalized Fung strain energy function (*W*) in Abaqus has the following form:

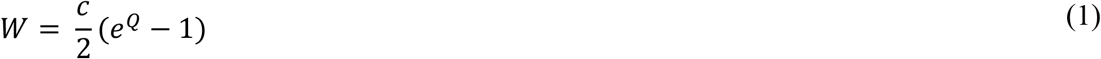

where *c* is the stress scaling factor and *Q* is given by

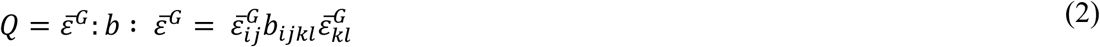

where *b*_*ijkl*_ is a dimensionless symmetric fourth-order tensor of anisotropic material parameters and 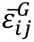 and 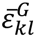 are the modified Green strain tensor. For the orthotropic case of the generalized Fung model, nine material parameters must be specified.

As explained by Sun and Sacks (2005), in the unloaded configuration, matrix *D* should be positive definite in order to obtain numerical stability. Matrix *D* is given by:

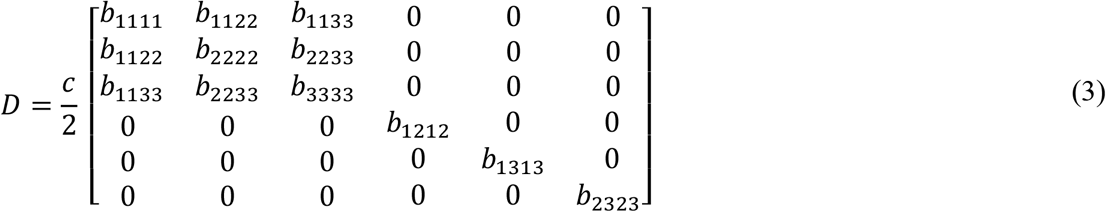

The positive definiteness implies the following parameter constraints:

- Constraint 1:

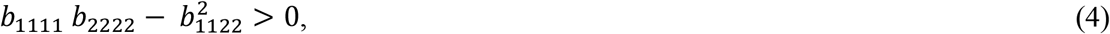
- Constraint 2:

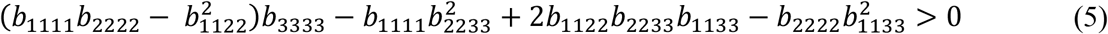
- Constraint 3:

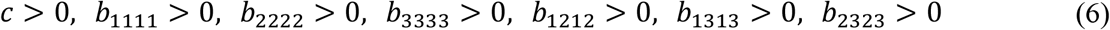

### 2.4 Optimisation of material parameters

The aim was to optimize the material parameters of Fung-orthotropic model to fit the FE model solution to the experimental stress-strain data for up to 5% of biaxial strain; the average physiological range of infarct deformation (Fomovsky and Holmes, 2010). Displacement-controlled FE simulations were run and the computed stress data were compared to the experimental data. The objective function (OBJ) utilized in the optimisation was developed so that it returns the mean absolute percentage error (MAPE). The GA toolbox in SCILAB 5.4 (Scilab Enterprises S.A.S, Versailles, France) was utilized to minimize the objective function. For each infarct model, the FE stress data were obtained from three simulations: M6060, M3060 and M6030, representing the biaxial loading protocols of 60:60, 30:60 and 60:30 N/m, respectively, applied in the experiment (Sirry et al., 2016a).

The objective function is given by:

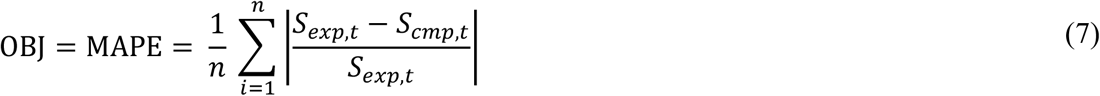

where *S*_*exp*_ and *S*_*cmp*_ are the experimental and computational stresses, respectively, *i* denotes the data point and *n* is the total number of fitted data points. In the optimisation process, *n* = 36 data points were utilized resulting from six equidistant points in each direction from three loading protocols. Upper and lower bounds were set for the material parameters as illustrated in Table 2. These bounds enforced the parameter constraints described in Eq.6.

The flow chart of the optimisation loops is shown in Figure 2. Initially, several sets of parameters (population) were randomly generated. A set (member) contained an array of 10 values representing the material parameters. A member would undergo a series of tests to evaluate the violation of Constraint 1 (Eq.4) and Constraint 2 (Eq.5). If a violation was detected, the member would be penalized by assigning an OBJ value according to the severity of the violation. Otherwise, the member would be assigned to Fung material parameters in Abaqus to start the FE simulations.

**Figure 2:**
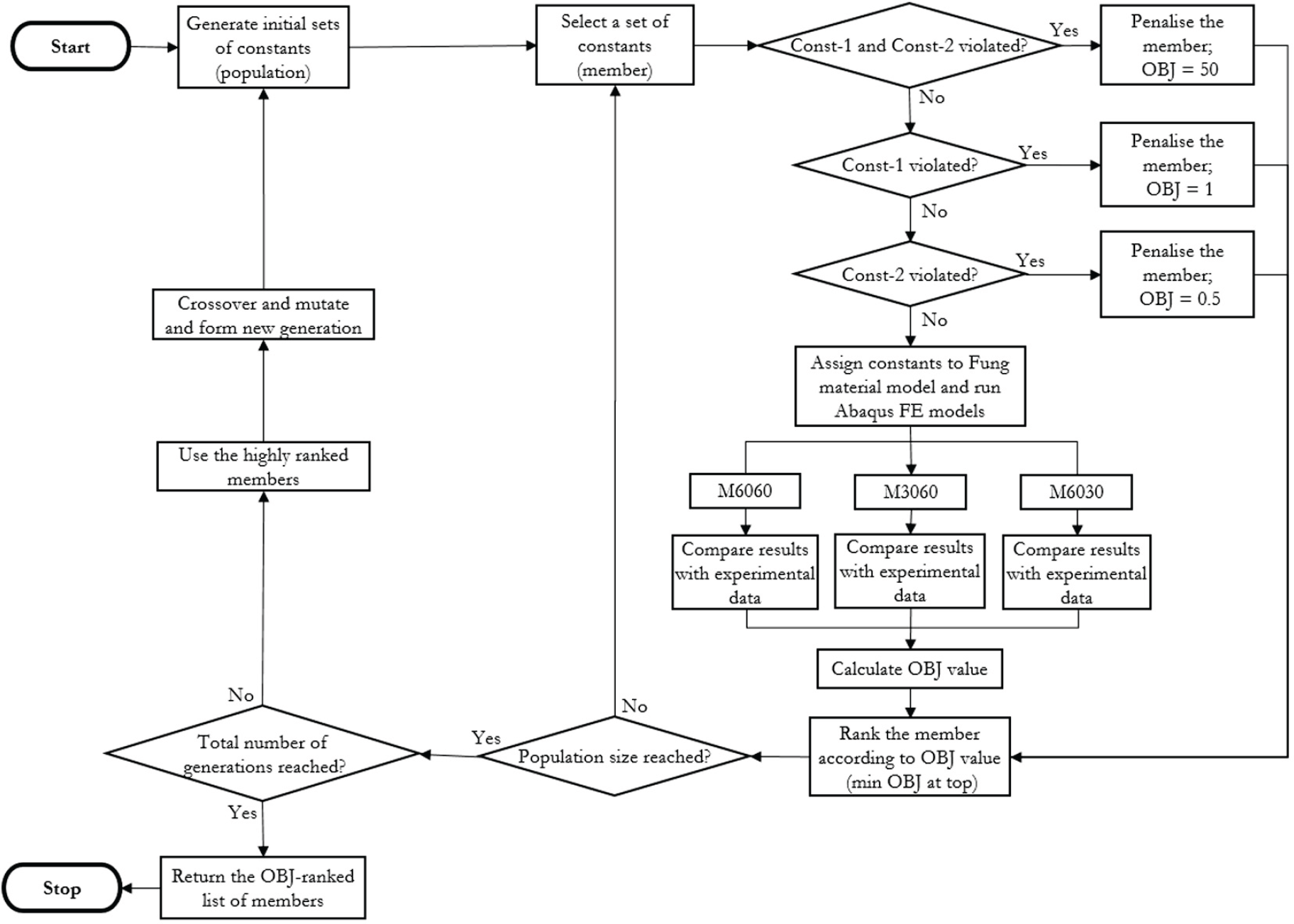
Flow chart of the SCILAB script utilized for identification of material parameters. Const-1 = Constraint 1, Const-2 = Constraint 2. OBJ = objective function.

At completion of FE simulations, computational and experimental stresses were compared and the OBJ value was calculated. After that, the member was ranked in such a way that the lower the OBJ value was, the higher the rank would be. The next member in the population underwent the same process until all members were assigned OBJ value and ranked. Then, a new generation was formed. This was accomplished by subjecting the highly ranked members to crossover (exchange of parameter values of two members) and mutation (changing parameters value in a member) processes to generate new members. The members of the new generation were ranked and a third generation was formed. The loop continued until the predefined number of generations was reached at which a list of the OBJ-ranked members was returned. The member with the smallest OBJ demonstrated the best fit. The GA predefined parameters are shown in Table 1.

**Table 1:** Predefined parameters used in set up of genetic algorithms.

| Parameter | Value | Note |
| --- | --- | --- |
| Population size | 100 | Number of members in a single generation (iteration) |
| Number of generations | 10 | Number of iterations |
| Number of couples | 50 | Number of member pairs (parents) in a population from which new members are formed through mutation and crossover |
| Mutation probability | 0.5 |  |
| Crossover probability | 0.7 |  |

### 2.5 Computation of stress and strain

The biaxial stress and strain were computed from the FE model in SCILAB environment. Python script was utilized to extract the required results from the model output-database file. The stress was computed based on the following equations:

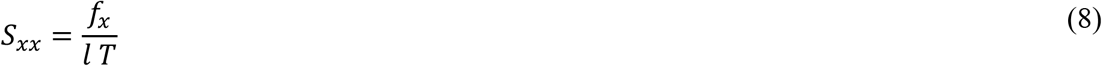

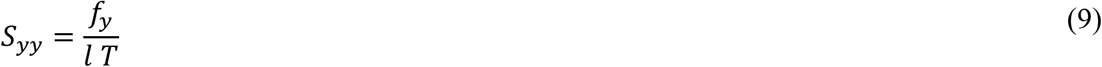

where *S*_*xx*_, *f*_*x*_ and *S*_*yy*_, *f*_*y*_ are engineering stresses and forces in the circumferential and longitudinal directions respectively, *l* and *T* are length and thickness of the model, respectively. The values of *f*_*x*_ and *f*_*y*_ were captured from the predicted reaction forces at C5 and C6, for *f*_*y*_, and C7 and C8, for *f*_*x*_, respectively (Figure 1).

The nodal strain was derived from the biaxial displacement of the four reference points (RP1-4 in Figure 1). Cartesian coordinates of these points were captured at each increment of the quasi-static simulation. The nodal strain was computed based on the following equations (Billiar and Sacks, 2000; Hoffman and Grigg, 1984; Humphrey et al., 1987):

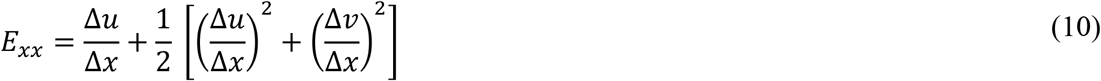

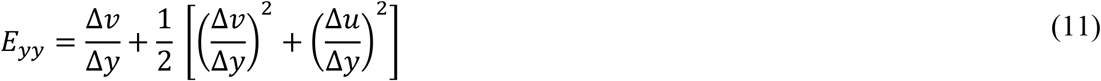

where *E*_*xx*_ and *E*_*yy*_ are the nodal strains, ∆*x* and ∆*y* are the changes in nodal coordinates, and ∆*u* and ∆*v* are the changes in nodal displacement in the circumferential and longitudinal directions, respectively.

## 3. Results

The identified Fung orthotropic material parameters for different infarct groups are shown in Table 2. The MAPE value represents the relative error between the experimental and computational data. The FE model predictions best approximated the experimental data of the 28d group with 3.0% error. The worst approximation was for the 7d group with 3.6% error. Comparisons between the experimental data and the best fitted FE model results are shown in Figure 3, Figure 4, Figure 5 and Figure 6 for the immediate, 7d, 14d and 28d infarct groups, respectively. Most of the numerical predictions fell within the standard deviation tolerance of the experimental data for different infarcts. Mechanical coupling exhibited by the experimental data, indicated by the negative circumferential strains, at 30:60 loading protocol was predicted by the FE model for all infarcts as illustrated in Figures 3-6 (c). With exception to the 28d group, FE models were not able to closely approximate the circumferential stress-strain data of the 60:30 protocol, Figures 3-6 (e).

**Figure 3:**
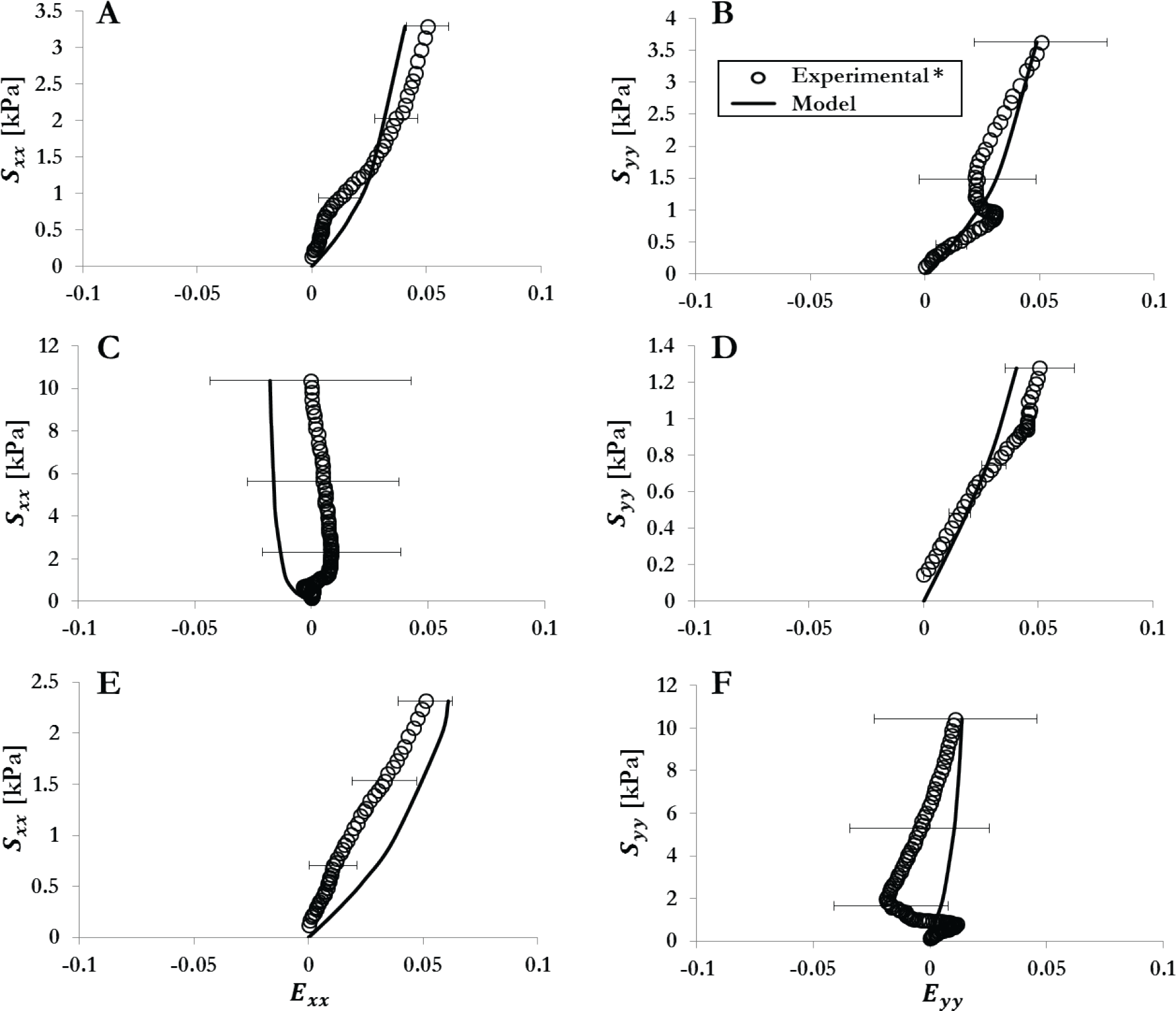
Best fit of FE model predictions to the experimental data for the immediate (0d) infarct group in each direction from three loading protocol. Stress axes are shown at variable range. Error bars represent the standard deviation. (a) Circumferential direction from 60:60 N/m protocol. (b) Longitudinal direction from 60:60 N/m protocol. (c) Circumferential direction from 30:60 N/m protocol. (d) Longitudinal direction from 30:60 N/m protocol. (e) Circumferential direction from 60:30 N/m protocol. (f) Longitudinal direction from 60:30 N/m protocol. * Experimental data obtained from Sirry et al. (Sirry et al., 2016b).

**Figure 4:**
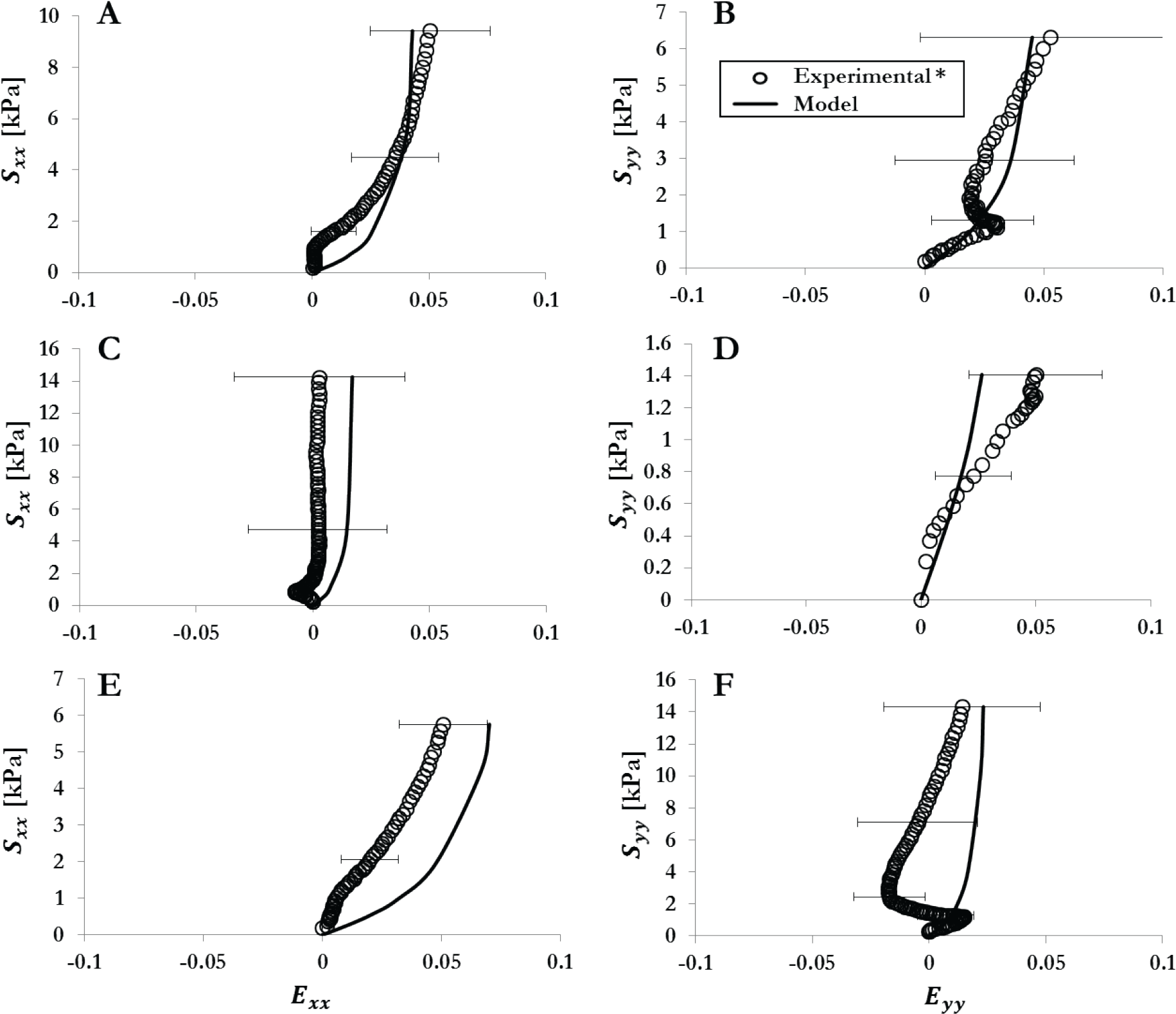
Best fit of FE model predictions to the experimental data for the 7d infarct group in each direction from three loading protocol. Stress axes are shown at variable range. Error bars represent the standard deviation. (a) Circumferential direction from 60:60 N/m protocol. (b) Longitudinal direction from 60:60 N/m protocol. (c) Circumferential direction from 30:60 N/m protocol. (d) Longitudinal direction from 30:60 N/m protocol. (e) Circumferential direction from 60:30 N/m protocol. (f) Longitudinal direction from 60:30 N/m protocol. * Experimental data obtained from Sirry et al. (Sirry et al., 2016b).

**Figure 5:**
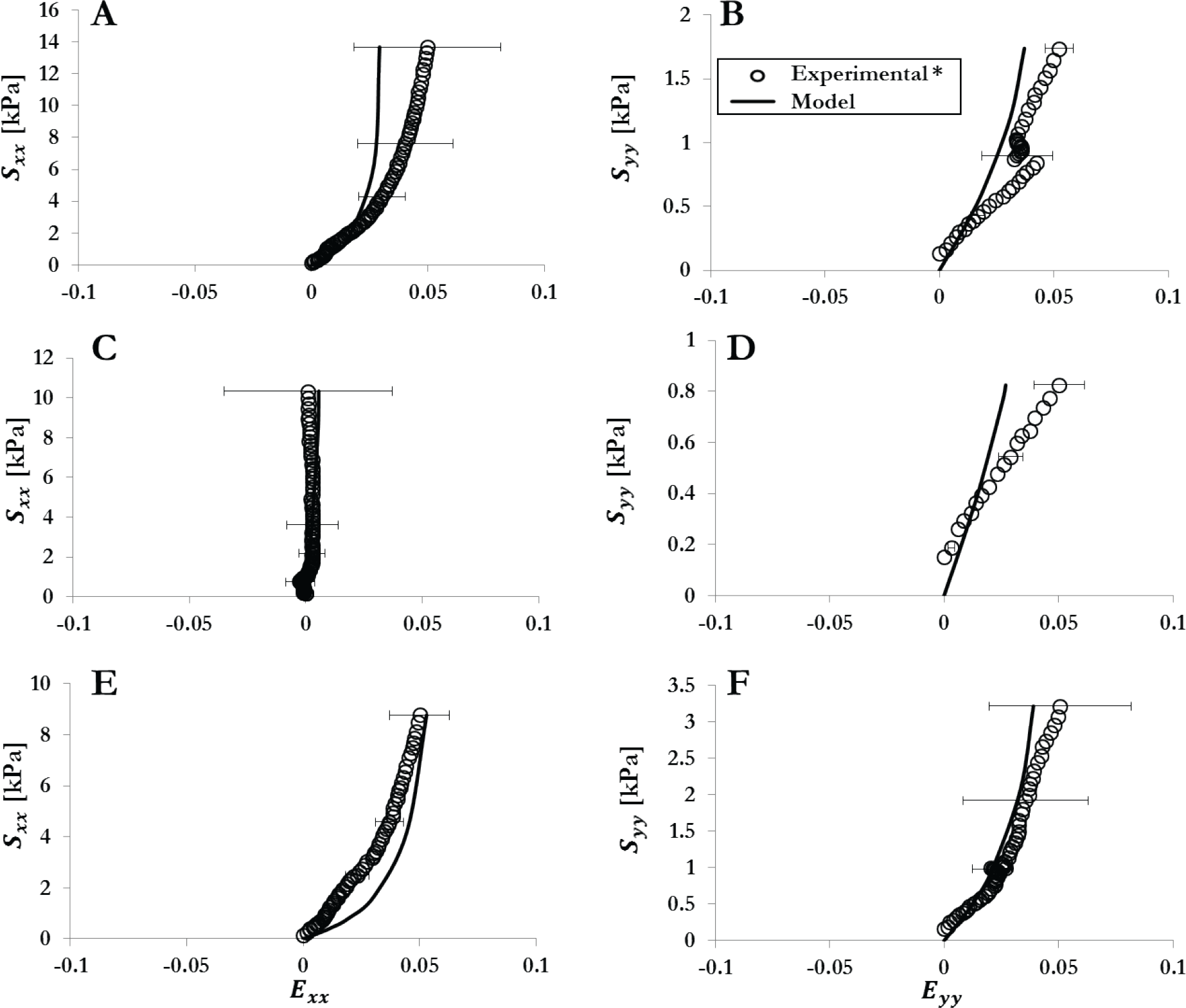
Best fit of FE model predictions to the experimental data for the 14d infarct group in each direction from three loading protocol. Stress axes are shown at variable range. Error bars represent the standard deviation. (a) Circumferential direction from 60:60 N/m protocol. (b) Longitudinal direction from 60:60 N/m protocol. (c) Circumferential direction from 30:60 N/m protocol. (d) Longitudinal direction from 30:60 N/m protocol. (e) Circumferential direction from 60:30 N/m protocol. (f) Longitudinal direction from 60:30 N/m protocol. * Experimental data obtained from (Sirry et al., 2016b).

**Figure 6:**
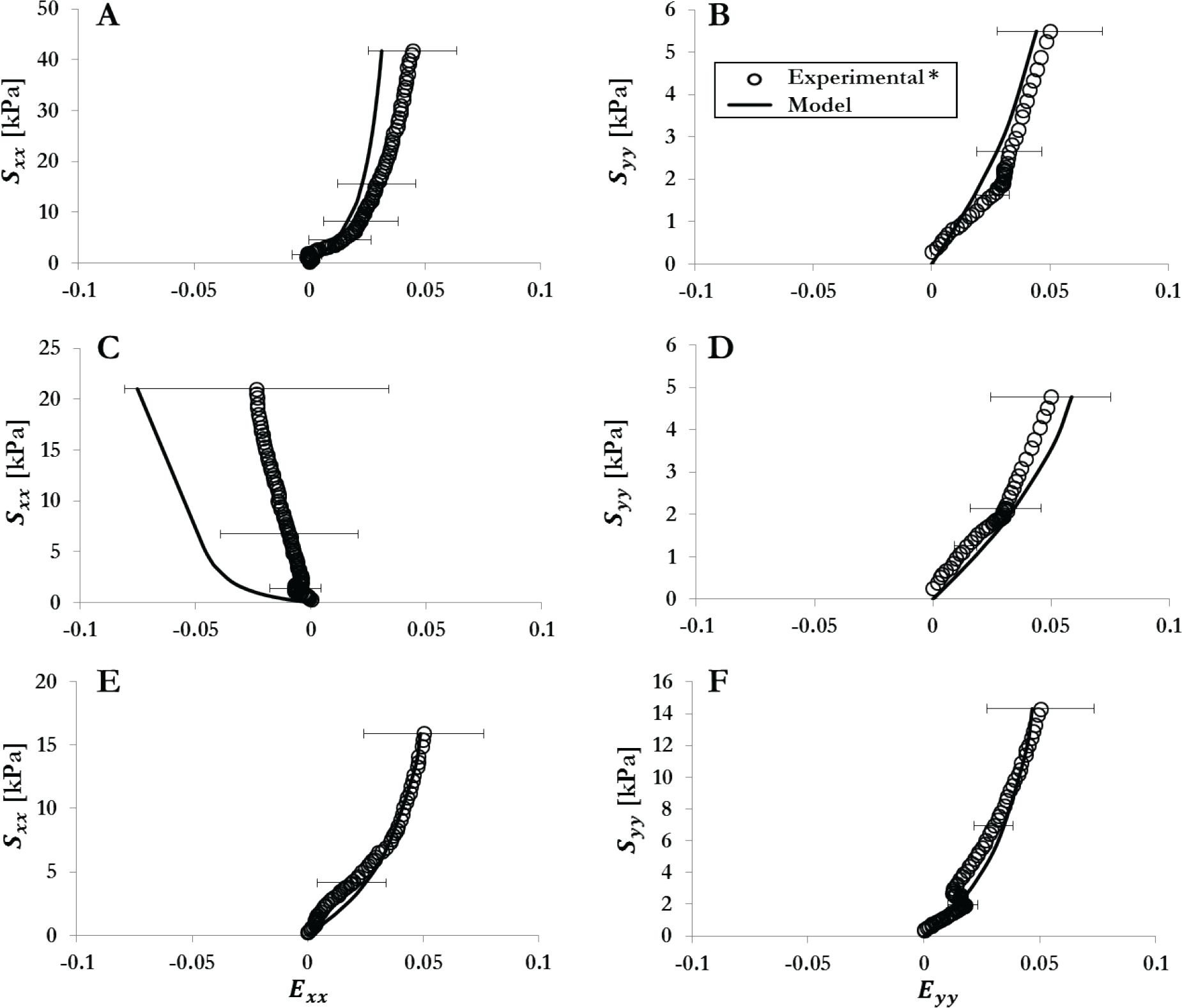
Best fit of FE model predictions to the experimental data for the 28d infarct group in each direction from three loading protocol. Stress axes are shown at variable range. Error bars represent the standard deviation. (a) Circumferential direction from 60:60 N/m protocol. (b) Longitudinal direction from 60:60 N/m protocol. (c) Circumferential direction from 30:60 N/m protocol. (d) Longitudinal direction from 30:60 N/m protocol. (e) Circumferential direction from 60:30 N/m protocol. (f) Longitudinal direction from 60:30 N/m protocol. * Experimental data obtained from (Sirry et al., 2016b).

## 4. Discussion

In this study, an inverse approach was applied to identify material parameters of the orthotropic case of Fung constitutive law for healing rat myocardial infarcts. Utilizing the experimental biaxial tensile data (Sirry et al., 2016b) from previous work (Sirry et al., 2016a), FE models were developed and numerical solutions were fitted to the experimental data through GA-based optimisation of the material parameters. We believe the material parameters identified in this study will provide a new tool for FE investigation of MI mechanics based on rat models. Stretching the model using 2 cylinders in each direction while constraining the movement of the 2 opposite cylinders produced uniform stress field in the central target area as illustrated in Figure 7. The locations of the reference points (black dots) were far enough from the areas of high stress concentration near the needles. This confirms that the model did not predict false strains due to the improper deformation of the sample (Sun et al., 2005).

**Figure 7:**
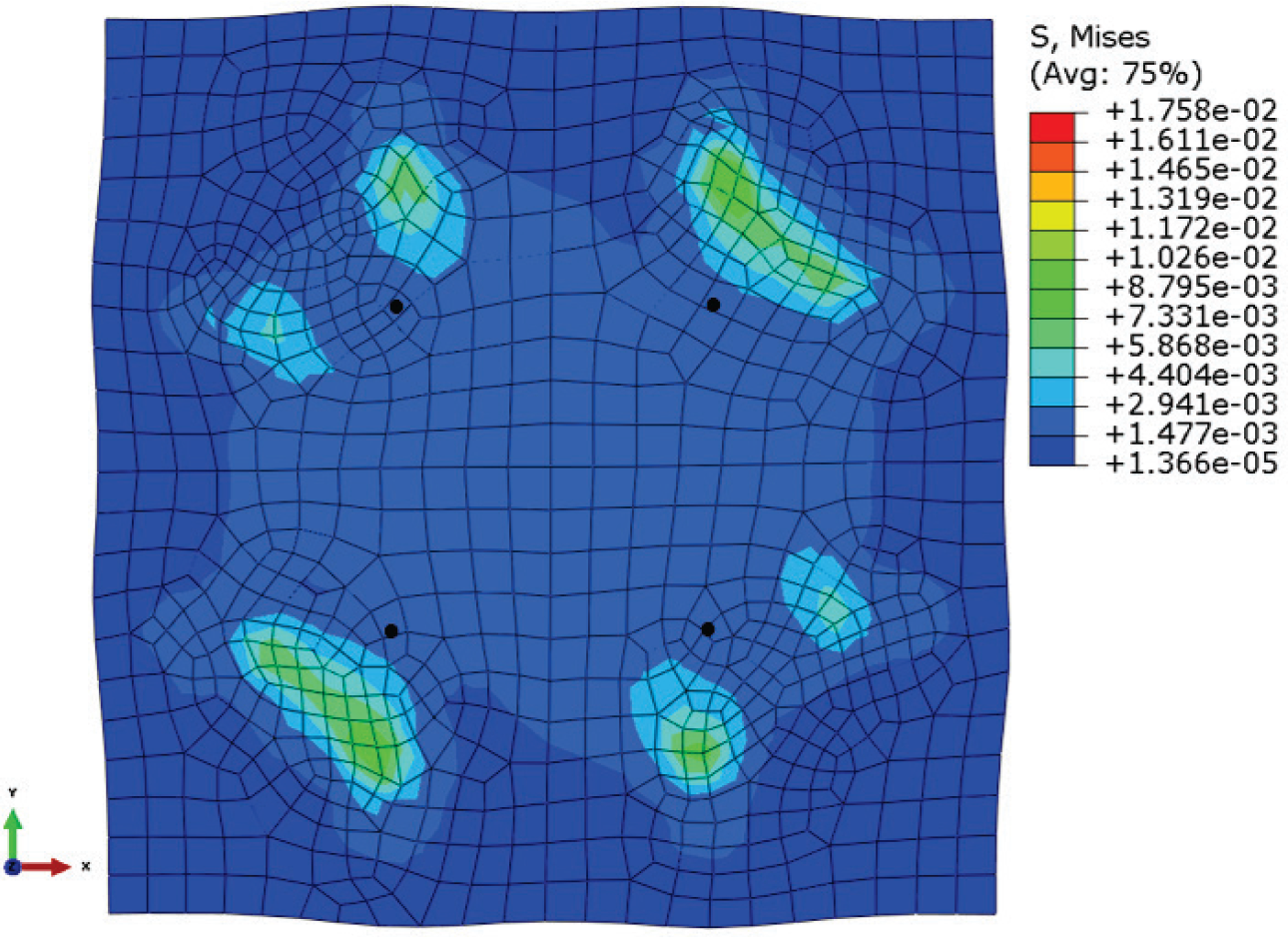
Von Mises stress (MPa) distribution in a fully deformed biaxial FE model illustrating the uniform stress field in the central target area. Locations of reference points are indicated by black dots.

It has been discussed that myocardium may have orthotropic structure and that orthotropic constitutive models better describe the mechanical behaviour of myocardium than transversely anisotropic models (Costa et al., 2001; LeGrice et al., 2001; Usyk et al., 2000). In the present study, an orthotropic constitutive law was used to model the mechanical properties of the different infarct stages.

One difficulty in optimizing the material parameters in this model was the positive definiteness condition of the parameter matrix (Eq.3) in the unloaded state. To fulfilment the constraints described in Eqs.4 and 5, it was verified that the generated sets of parameters satisfied the positive definiteness condition, similar to Wang et al. (2006). In a generation, random sets of material parameters were defined by the GA. It was required to test these sets for violation of the constraints before they were passed to the model. The set that violates Constraints 1 and 2 was the worst and penalized using a relatively large OBJ value (OBJ=50). If only Const-1 was violated, the set was penalized with less OBJ value (OBJ=1). If only Constraint 2 was violated, the set was penalized with OBJ=0.5. Penalizing the undesired sets would rank them at the bottom of the list so that they were not included in the formation of the next generation. This way the GA were biased in the definition of the new generation. As a result, the violation disappeared in most of the newly formed sets of parameters at the 4^th^ or 5^th^ generation.

A shortcoming of the current study was the employment of biaxial data only to fit material parameters of orthotropic material law. The negative values in some of the identified parameters, e.g. *b*_1122_ and *b*_2233_ (Table 2), did not make good physical sense and indicated “overfitting”. In other words, the biaxial tests were not able to provide sufficient data to fit all the parameters of the orthotropic constitutive law. Holzapfel and Ogden (2009) discussed the limitation of using biaxial data alone to fit orthotropic constitutive law. They emphasized the need for more comprehensive mechanical characterization which include shear data at different orientation in order to appropriately capture the orthotropic behaviour of myocardial tissue. Nevertheless, the parameters identified in this study would still be valuable establishment.

**Table 2:**
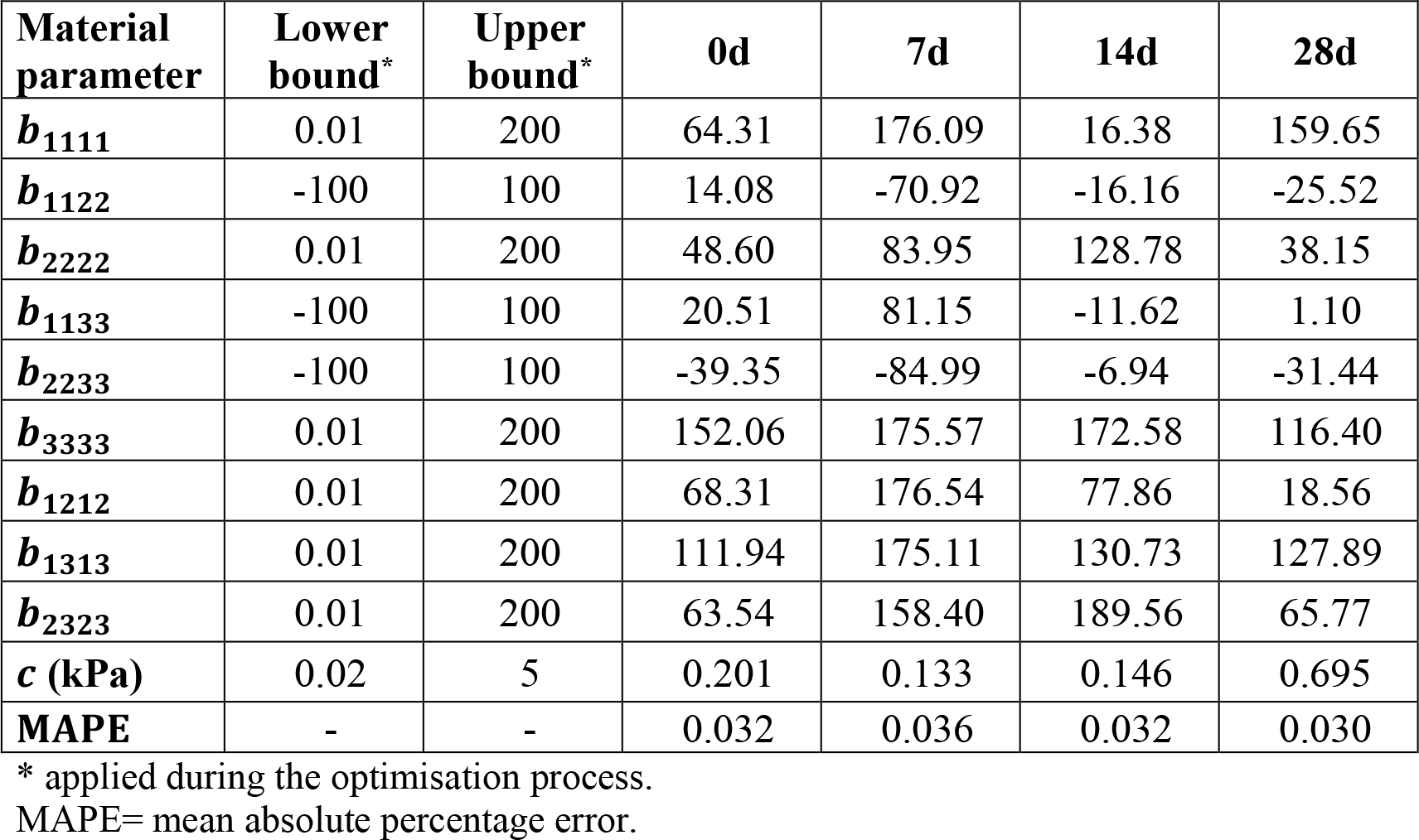
Identified Fung orthotropic material parameters for immediate, 7, 14 and 28 day infarcts.

| **Material parameter** | **Lower bound\*** | **Upper bound\*** | **0d** | **7d** | **14d** | **28d** |
| --- | --- | --- | --- | --- | --- | --- |
| **$b_{1111}$** | 0.01 | 200 | 64.31 | 176.09 | 16.38 | 159.65 |
| **$b_{1122}$** | -100 | 100 | 14.08 | -70.92 | -16.16 | -25.52 |
| **$b_{2222}$** | 0.01 | 200 | 48.60 | 83.95 | 128.78 | 38.15 |
| **$b_{1133}$** | -100 | 100 | 20.51 | 81.15 | -11.62 | 1.10 |
| **$b_{2233}$** | -100 | 100 | -39.35 | -84.99 | -6.94 | -31.44 |
| **$b_{3333}$** | 0.01 | 200 | 152.06 | 175.57 | 172.58 | 116.40 |
| **$b_{1212}$** | 0.01 | 200 | 68.31 | 176.54 | 77.86 | 18.56 |
| **$b_{1313}$** | 0.01 | 200 | 111.94 | 175.11 | 130.73 | 127.89 |
| **$b_{2323}$** | 0.01 | 200 | 63.54 | 158.40 | 189.56 | 65.77 |
| **$c$ (kPa)** | 0.02 | 5 | 0.201 | 0.133 | 0.146 | 0.695 |
| **MAPE** | - | - | 0.032 | 0.036 | 0.032 | 0.030 |

\* applied during the optimisation process.
MAPE= mean absolute percentage error.

## 5. Conclusions

We successfully employed the biaxial mechanical data to establish anisotropic constitutive models of healing infarcts in the rat model. Inverse finite element model and genetic algorithm were employed to identify the parameters of Fung orthotropic hyperelastic strain energy function for the healing infarcted tissue. The identified material parameters will provide new platform for FE investigations of mechanical aspects of MI and therapies in particular when using rat models.

## Acknowledgements

This study was supported financially by the National Research Foundation (NRF) of South Africa (UID92531) and the Centre for High Performance Computing, Council for Scientific and Industrial Research, South Africa (UCTUO18697). Any opinion, findings and conclusions or recommendations expressed in this publication are those of the authors and therefore the NRF does not accept any liability in regard thereto. MSS acknowledges the International Society of Biomechanics Matching Dissertation Grant. NHD acknowledges financial support from the South African Medical Research Council.

## Data

Data used in this study are available on ZivaHUB: http://doi.org/10.25375/uct.9757577.

Data include:

- Mean stress-strain data from biaxial tensile tests of myocardial samples [Stress-strain data from biaxial tests of individual samples are available from Sirry et al. (2016b)]
- Abaqus input files of FE models
- Optimization algorithms

## References

Billiar, K.L., Sacks, M.S., 2000. Biaxial mechanical properties of the natural and glutaraldehyde treated aortic valve cusp--part i: Experimental results. Journal of Biomechanical Engineering 122, 23–30.

Chen, J., Song, S.-K., Liu, W., McLean, M., Allen, J.S., Tan, J., Wickline, S.A., Yu, X., 2003. Remodeling of cardiac fiber structure after infarction in rats quantified with diffusion tensor mri. American Journal of Physiology – Heart and Circulatory Physiology 285, H946–H954.

Costa, K.D., Holmes, J.W., McCulloch, A.D., 2001. Modelling cardiac mechanical properties in three dimensions. Royal Society of London Philosophical Transactions Series A 359, 1233–1250.

Fomovsky, G.M., Holmes, J.W., 2010. Evolution of scar structure, mechanics, and ventricular function after myocardial infarction in the rat. American Journal of Physiology – Heart and Circulatory Physiology 298, H221–H228.

Fung, Y.C., Fronek, K., Patitucci, P., 1979. Pseudoelasticity of arteries and the choice of its mathematical expression. American Journal of Physiology 237, H620–H631.

Guccione, J.M., McCulloch, A.D., Waldman, L.K., 1991. Passive material properties of intact ventricular myocardium determined from a cylindrical model. Journal of Biomechanical Engineering 113, 42–55.

Gupta, K.B., Ratcliffe, M.B., Fallert, M.A., Edmunds, L.H., Bogen, D.K., 1994. Changes in passive mechanical stiffness of myocardial tissue with aneurysm formation. Circulation 89, 2315–2326.

Hoffman, A.H., Grigg, P., 1984. A method for measuring strains in soft tissue. Journal of Biomechanics 17, 795–800.

Holzapfel, G.A., Ogden, R.W., 2009. Constitutive modelling of passive myocardium: A structurally based framework for material characterization. Philosophical Transactions of the Royal Society a-Mathematical Physical and Engineering Sciences 367, 3445–3475.

Humphrey, J.D., 1995. Mechanics of the arterial wall: Review and directions. Critical Reviews in Biomedical Engineering 23, 1–162.

Humphrey, J.D., Vawter, D.L., Vito, R.P., 1987. Quantification of strains in biaxially tested soft tissues. Journal of Biomechanics 20, 59–65.

Humphrey, J.D., Strumpf, R.K., Yin, F.C.P., 1990. Determination of a constitutive relation for passive myocardium: Ii.--- parameter estimation. Journal of Biomechanical Engineering 112, 340–346.

Khalil, A., Bouma, B., Kaazempur Mofrad, M., 2006. A combined fem/genetic algorithm for vascular soft tissue elasticity estimation. Cardiovascular Engineering 6, 93–102.

Kichula, E., Wang, H., Dorsey, S., Szczesny, S., Elliott, D., Burdick, J., Wenk, J., 2014. Experimental and computational investigation of altered mechanical properties in myocardium after hydrogel injection. Annals of Biomedical Engineering 42, 1546–1556.

Kortsmit, J., Davies, N.H., Miller, R., Macadangdang, J.R., Zilla, P., Franz, T., 2013. The effect of hydrogel injection on cardiac function and myocardial mechanics in a computational post-infarction model. Computer Methods in Biomechanics and Biomedical Engineering 16, 1185–1195.

Lee, L.C., Wenk, J.F., Klepach, D., Zhang, Z., Saloner, D., Wallace, A.W., Ge, L., Ratcliffe, M.B., Guccione, J.M., 2011. A novel method for quantifying in-vivo regional left ventricular myocardial contractility in the border zone of a myocardial infarction. Journal of Biomechanical Engineering 133, 094506–094505.

LeGrice, I., Hunter, P., Young, A., Smaill, B., 2001. The architecture of the heart: A data-based model. Philosophical Transactions of the Royal Society A: Mathematical, Physical and Engineering Sciences 359, 1217–1232.

Miller, R., Davies, N.H., Kortsmit, J., Zilla, P., Franz, T., 2013. Outcomes of myocardial infarction hydrogel injection therapy in the human left ventricle dependent on injectate distribution. International Journal for Numerical Methods in Biomedical Engineering 29, 870–884.

Sirry, M.S., Butler, J.R., Patnaik, S.S., Brazile, B., Bertucci, R., Claude, A., McLaughlin, R., Davies, N.H., Liao, J., Franz, T., 2016a. Characterisation of the mechanical properties of infarcted myocardium in the rat under biaxial tension and uniaxial compression. Journal of the Mechanical Behavior of Biomedical Materials 63, 252–264.

Sirry, M.S., Butler, J.R., Patnaik, S.S., Brazile, B., Bertucci, R., Claude, A., McLaughlin, R., Davies, N.H., Liao, J., Franz, T., 2016b. Infarcted rat myocardium: Data from biaxial tensile and uniaxial compressive testing and analysis of collagen fibre orientation. Data in Brief 8, 1338–1343.

Sun, W., Sacks, M.S., 2005. Finite element implementation of a generalized fung-elastic constitutive model for planar soft tissues. Biomechanics and Modeling in Mechanobiology 4, 190–199.

Sun, W., Sacks, M.S., Scott, M.J., 2005. Effects of boundary conditions on the estimation of the planar biaxial mechanical properties of soft tissues. Journal of Biomechanical Engineering 127, 709–715.

Usyk, T.P., McCulloch, A.D., Computational methods for soft tissue biomechanics., in: Holzapfel, G. A. Ogden, R. W., Eds.), Biomechanics of soft tissue in cardiovascular systems., Springer, Wien 2003, pp. 273–342.

Usyk, T.P., Mazhari, R., McCulloch, A.D., 2000. Effect of laminar orthotropic myofiber architecture on regional stress and strain in the canine left ventricle. Journal of Elasticity 61, 143–164.

Wang, C., Garcia, M., Lu, X., Lanir, Y., Kassab, G.S., 2006. Three-dimensional mechanical properties of porcine coronary arteries: A validated two-layer model.

Wenk, J., Zhang, Z., Cheng, G., Malhotra, D., Acevedo-Bolton, G., Burger, M., Suzuki, T., Saloner, D., Wallace, A., Guccione, J., 2010. First finite element model of the left ventricle with mitral valve: Insights into ischemic mitral regurgitation. The Annals of Thoracic Surgery 89, 1546–1553.

Wenk, J.F., Sun, K., Zhang, Z., Soleimani, M., Ge, L., Saloner, D., Wallace, A.W., Ratcliffe, M.B., Guccione, J.M., 2011. Regional left ventricular myocardial contractility and stress in a finite element model of posterobasal myocardial infarction. Journal of Biomechanical Engineering 133, 044501.

Wise, P., Davies, N.H., Sirry, M.S., Kortsmit, J., Dubuis, L., Chai, C.-K., Baaijens, F.P.T., Franz, T., 2016. Excessive volume of hydrogel injectates may compromise the efficacy for the treatment of acute myocardial infarction. International Journal for Numerical Methods in Biomedical Engineering.

Yeoman, M.S., Reddy, B.D., Bowles, H.C., Zilla, P., Bezuidenhout, D., Franz, T., 2009. The use of finite element methods and genetic algorithms in search of an optimal fabric reinforced porous graft system. Annals of Biomedical Engineering 37, 2266–2287.

